# Land Use Land Cover and Change Detection Analysis in Singareni Opencast Coalmines Area of Telangana State using RS and GIS Technique

**DOI:** 10.64898/2026.08.07.743447

**Authors:** J Kamalakar, T Ram Prakash, S Jhonson Raju, G Jayashree, M. Yakadri

## Abstract

Investigating Changes Detection in Land Use and Land Cover Analysis of Opencast Singareni Coal Mine Area Using Remote Sensing and GIS Techniques was the focus of the current investigation. Change Detection Analysis for Land Use Land Cover over a decade frequency (2005–2015) of cultivated soils surrounding OpenCast Coal Mine areas namely Ramakrishnapur, Srirampur and Medipalli of Telangana State.AWiFS data of IRS Resoursesat Satellite images of 2005 and 2015 were taken for the detection of temporal variation in Land use/ Land cover due to open cast coal mining.The ability of remote sensing techniques to produce precise spatiotemporal statistics of LULC and its changes in the typical coal mining region has been demonstrated, in open cast coal mining per year was the highest at Ramakrishnapur (@ 300 ha / year) followed by Srirampur (@ 150 ha / year) and Medipalli (@ 100 ha / year).At Ramakrishnapur with the onset of mining there was considerable decrease in agricultural land, shallow water bodies and deciduous forests *i.e.,* conversion of these land uses to mining was evident. Whereas, at other two mining sites (Srirampur and Medipalli) conversion of scrubland followed by shallow water bodies, agriculture lands and deciduous forests into mining area was detected. However, the built-up land at all three-mining site was from cultivated lands and at Medipalli and Srirampur it was also contributed from scrubland. On the whole impact of open cast mining on deep water bodies was not detected. If the mining and surrounding regions’ continued and appropriate use of the land and water resources at hand, careful long-term planning is of utmost importance.

## Introduction

There is a requirement to comprehend the dynamic surface and structural changes in a landscape, which can be successfully tracked using remote sensing and Geographic Information System approaches (1,2,3). GIS is a sophisticated tool for presenting and analyzing spatial relationships between geographic phenomena using vectors and pictures (3,4). Thematic mapping has undergone a revolution in recent years as a result of improvements in the geographic information system (1). Integration of Remote Sensing into a GIS database can minimize costs, shorten survey times, and enhance the amount of comprehensive information collected (5). There is still significant untapped potential in Indian soils to increase crop output through efficient and sustainable use of soil resources, including water and nutrients (6,7).

Land use and land cover (LU/LC) can be classified via remote sensing region’s quantitative digital image processing techniques or qualitative visual interpretation approaches (8,1,9). Even with quantitative approaches based on objective numerical methods, classification outputs are usually scene-dependent, making it difficult to apply the same processing strategy to an image scene with varying properties with the same level of success (10,11,12). Long-term LU/LC classification and change detection systems based on remote sensing face difficulty as the dynamics of LULC change in an active mining area naturally affect the picture sceneries (13,14,10). Another issue is the availability of data for the same season across numerous years. As a result of the seasonal variation, the LU/LC Classification and Change Analysis may contain additional inaccuracies (12). Despite these disadvantages, a number of employees have successfully tracked coal mining activities and their impacts on the environment, terrain, and LU/LC over very long time periods using remote sensing photos. (8,16,17,18,19).

The current study offered an approach to incorporating landscape ecology principles into land use plans, with the goal of reducing the effect of land conversion by guiding such transformations in ecology, including proper edaphic direction (20,21,22). Furthermore, the study aims to comprehend, develop, and demonstrate a methodology for an integrated application of Remote Sensing (RS), Geographic Information System (GIS), Landscape ecological principles in anticipating Land Use, Land Cover changes (LU/LC) due to open cast coal mining and in developing rehabilitation strategies based on the principle of prioritization in the open cast mining areas of Singareni Coal Mine area, in the Telangana region.

## MATERIALS AND METHODS

### Change Detection

Satellite images of the research area were acquired from 2005 to 2015 for this investigation. Awifs data for 2005 and 2015 were provided by the National Remote Sensing Centre (NRSC) in Hyderabad (23). A topographical map of the region obtained from the Indian survey was used for ground reference (Fig 1). The topographical map of India from the Survey of India (SOI) was rectified using geographic latitude-longitude and WGS 84 datum. A satellite image was registered using a georeferenced topographical map and both a map to image and image to image registration technique (24,3). The ERDAS imagine programme version 9.2 was used for georeferencing and registration (24). Table 1 details the satellite and auxiliary data used.

**Figure 1.**
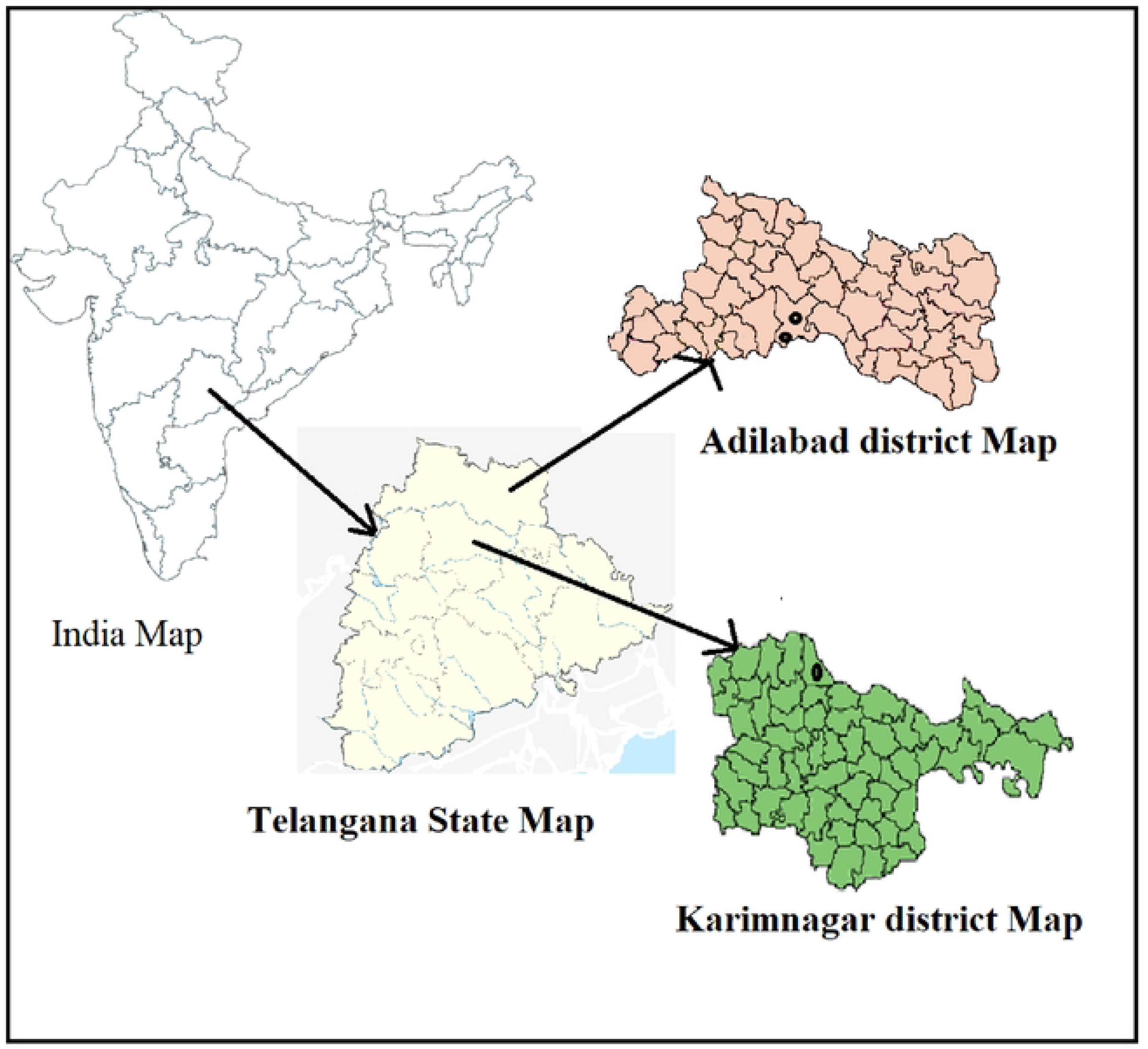
Location map of the study Area

**Table 1.**
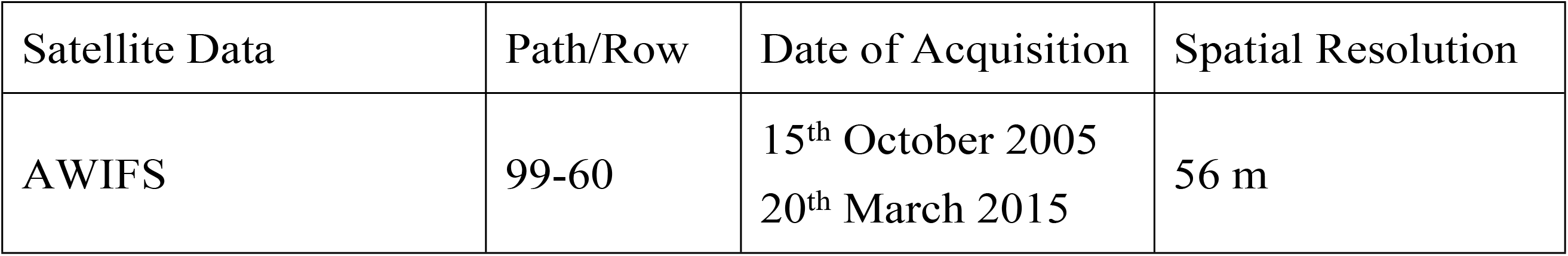
List of data used and their specifications.

### Land Use Land Cover Map

Comparing maps and photographs can reveal information on changes in land use and land cover. The current study used map-to-map comparison to determine changes in land use and land cover (14,25). Temporal satellite pictures were used to construct land utilization and land cover maps. Image processing and visual interpretation techniques were used to track changes in land use and cover. Tone, texture, shape, size, association, drainage, landform, soil, and vegetation are among the photographic and geotechnical features used to identify and classify the various land use/land cover groups (1,2,26,27). Image interpretation of satellite data was done in 2015 with reference to mapped Land use - Landover categories, followed by field validation. Using satellite data assembled during 2005 and 2015, we identified spatiotemporal changes in the region’s LU/LC over time. The National Remote Sensing Agency’s LU/LC classification scheme served as the foundation for the LU/LC categorization used in this study (28,29). The level-II category changes were made with the area’s coal mining activity and related Land Use - Land Cover in mind. Built-up, Agricultural Land, Plantation / Orchard, Deciduous Forest, Scrub Land, Waterbodies, Shallow waterbodies / Wet land, and Mining region were the eight classes that made up the level-II categorization. Using Arc Map software (version 9.3), an on-screen digitization approach was used to digitize the maps, and additional area statistics for other land use categories were computed. Using land use-land cover information between 2005 and 2015, the percentage shift, pattern, and pace of LU/LC change were calculated. The area and percent change for each land use and land cover type over a ten-year period were calculated. Using the following calculation, we were able to compute the % change by comparing the original (before) and final (after) LU/LC areal coverage (30,31):

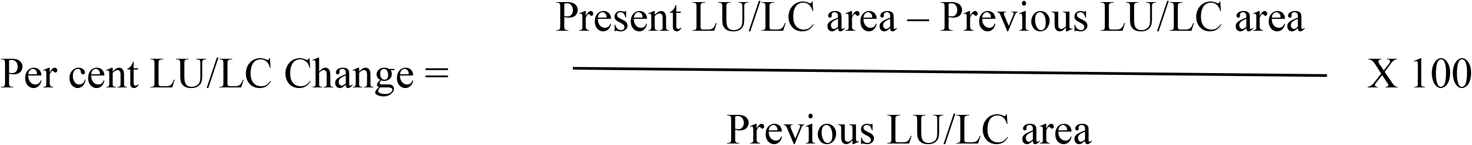

where area denotes the size of each LU/LC type. Positive values indicate expansion, whereas negative values indicate contraction. The methodology flow chart details the research approach employed. (Fig 5).

## RESULTS AND DISCUSSION

### Change detection of Land use / Land cover (LULC)

The land use of a region is the product of previous responses that projected the connections between the cultural, ecological, and physical contexts. As a result, a location’s land use pattern is the result of how people have utilised natural and socioeconomic components over time and space (32,33). An attempt was made to detect temporal variation of LU /LC over a period of 10 years *i.e.,* 2005 to 2015 with open cast coal mining in selected mining sites *i.e.,* Ramakrishnapur, Medipalli and Srirampur. AWiFS data of IRS Resoursesat satellite images of 2005 and Resoursesat 2015 were used for this purpose (23). As the pedons and surface samples were collected at different radial distances from opencast coal mining, the satellite image was also clipped with a radius of 10 km approximately from open cast coal mining for all three mining places separately and the images digitally processed under ERDAS 9.2 software (24,34) and the steps involved were shown in the flow chart (Fig 5).

The use of remote sensing in conjunction with a geographic information system (GIS) provides the most precise means of monitoring the extent and pattern of LU/LC changes in land scape conditions across time (1,2). The major land use/land cover categories in the area were mapped using satellite images from 2005 and 2015 (Figs 2, 3, and 4). Built-up land, agricultural area, plantation / orchards, shallow water bodies / wet land, deciduous forest, scrub land, water bodies, and mining area are the eight major categories.

**1. Built-up land** is territory that is covered by population-related settlements. The loss of farmland and grassland areas is often linked to urban development and the resulting population growth (32). Built land can be identified on false colour composite (FCC) by its cyan-light grey tone, coarse texture, dispersed pattern, and uneven shape.
**2. Agricultural land** can be summarised as land utilised primarily for the production of food and fibre. It includes agricultural land. (irrigated and non-irrigated, present fallow, and so on). Croplands are places with standing crops as of the date of the satellite above. Cropped regions are bright red to crimson in colour, have varying sizes and shapes, and have a regular or sub-regular outline shape in a contiguous to non-contiguous pattern. Fallow lands are cultivated fields that are temporarily un-cropped for one or more seasons, but not for less than one year or more than five years. On the FCC image, its light brown to light yellow tone, medium to smooth texture, non-contiguous pattern, and uneven form were used to identify it.
**3. Plantation/Orchard** regions are planted with agricultural tree crops using agricultural management techniques. These also include land use strategies and practices in which the cultivation of herbs, shrubs, and vegetable crops is purposely integrated with agricultural crops, generally in irrigation settings, for ecological and financial reasons. Plantations have irregular and sharp boundaries and are dark-red to crimson in colour, indicating that they are surrounded by a fence.
**4. A deciduous forest** is characterized as land having a tree canopy cover of more than 10% and an area greater than 0.5 hectares. Forests are determined both by the presence of trees and the absence of other predominant land uses within the notified forest boundaries (35,36). Dense Forest can be interpreted from the FCC image by its dark red tone, coarse - medium texture, contiguous pattern and regular to irregular shape. Open Forest exhibit crown density in between 40% to 10%. It is easily identified on FCC image by its light red - pinkish colour, smooth - medium texture, contiguous to noncontiguous pattern with irregular outline.
**5. Scrub land:** Scrub has shallow and skeletal soils that are sometimes chemically damaged, extreme slopes that are badly eroded, and lands that are prone to excessive aridity where scrubs dominate the landscape (37). They have a tendency for intermixing with cropped areas. On the FCC image it can be identified by its pink - light yellow tone, coarse to medium texture scattered pattern and irregular outline boundary. If the vegetation cover is more dense it is called as dense scrub and open scrub if it is having thin soil cover.
**6. Water Bodies** are areas containing surface water, either impounded as ponds, lakes, and reservoirs or flowing as streams, rivers, canals, and so on. Depending on the depth of the water, these can be seen clearly in blue to dark blue or cyan on the satellite view (38).
**7. Shallow Water Bodies / Wet Land** are places with very shallow water bodies. These lakes of water arise as a result of the collection of rainwater or ground water in mining depressions (39,40).
**8. Mining area** denoted by blue to dark tones in the satellite image with mottled texture nearby current fallow and forest area as shown in satellite images Figs 2, 3 and 4.

**Figure 2.**
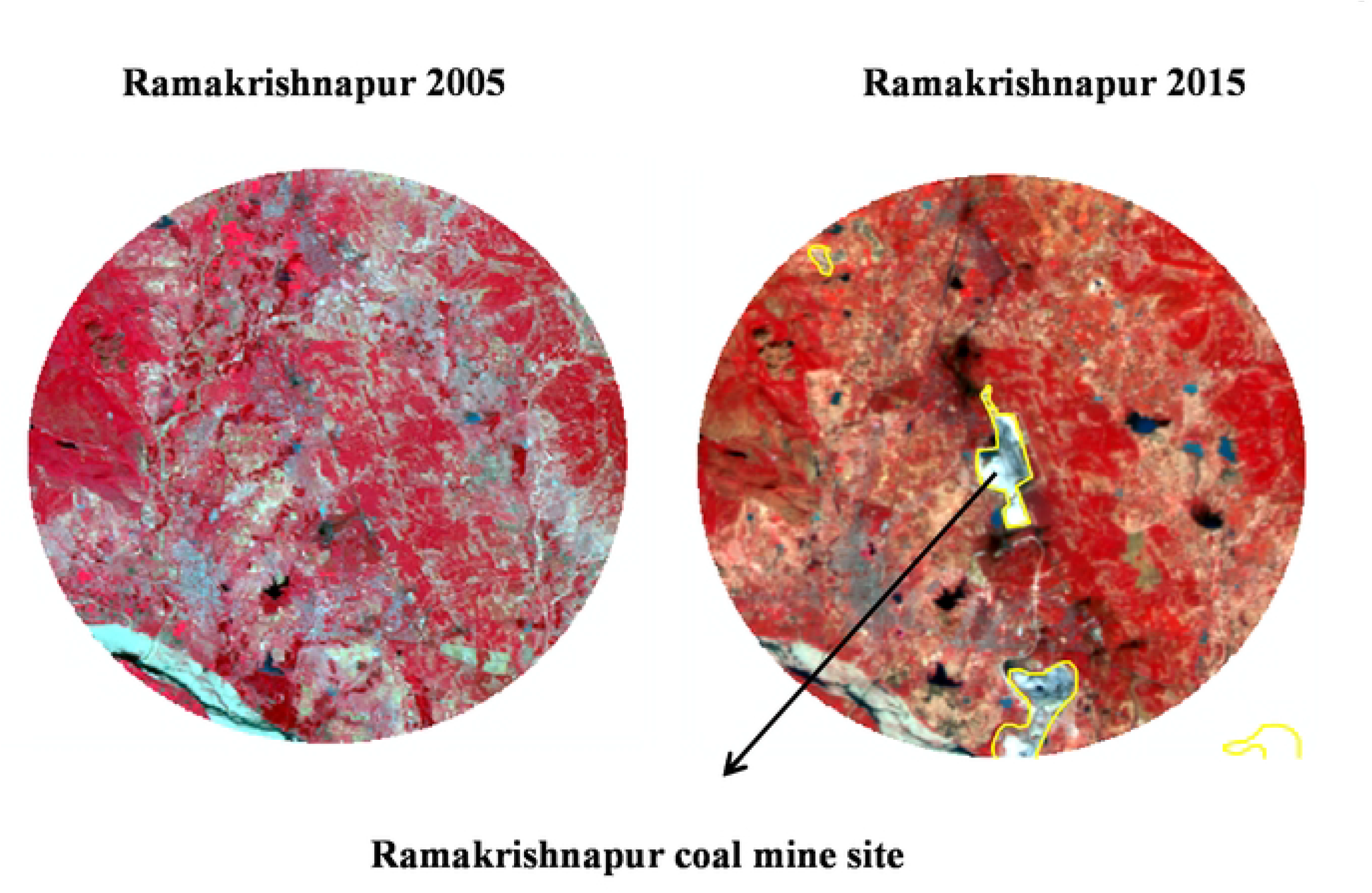
Satellite images of Ramakrishnapur open cast coal mine area a) Year 2005 b) Year 2015

**Figure 3.**
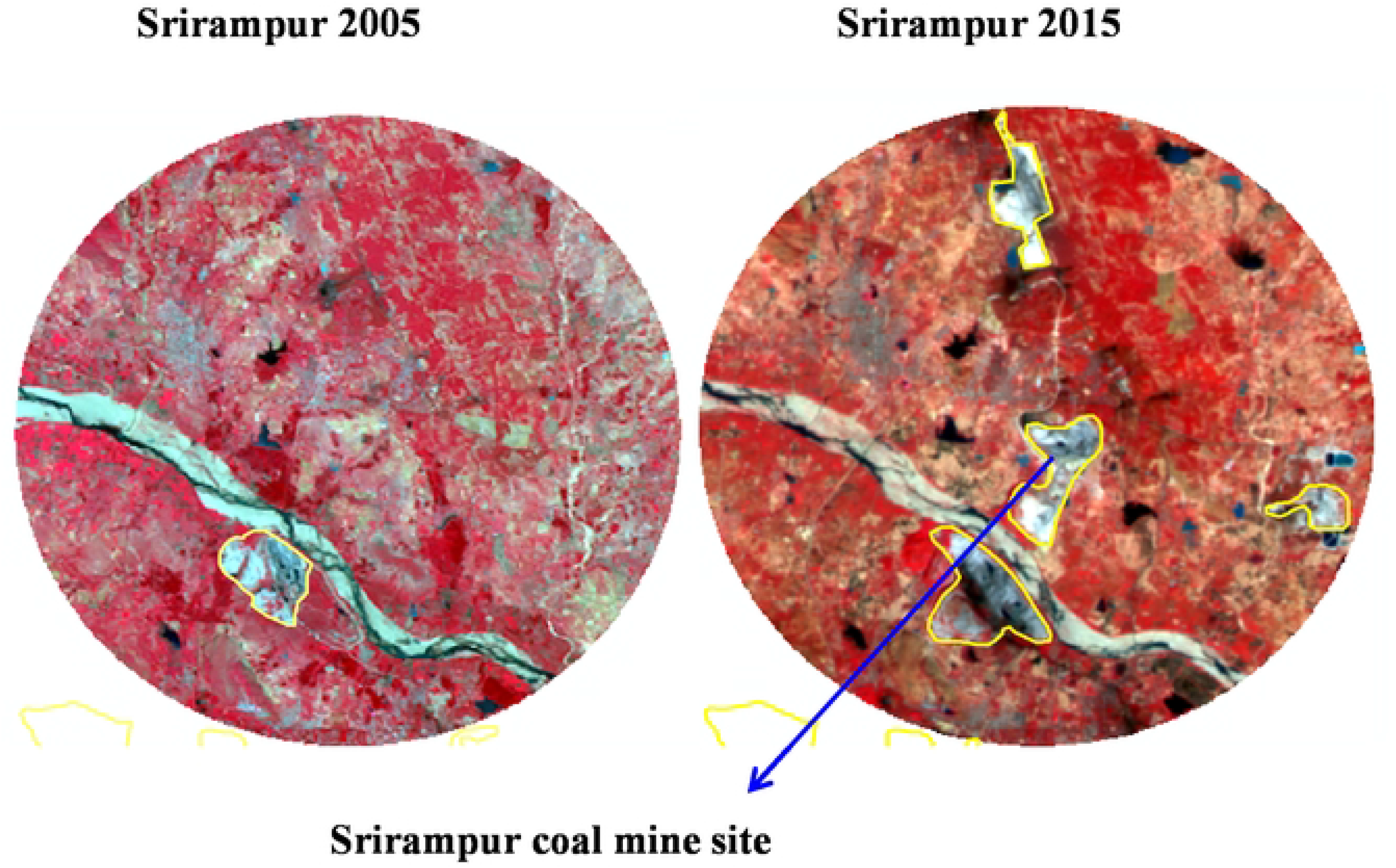
Satellite images of Srirampur open cast coal mine area a) Year 2005 b) Year 2015

**Figure 4.**
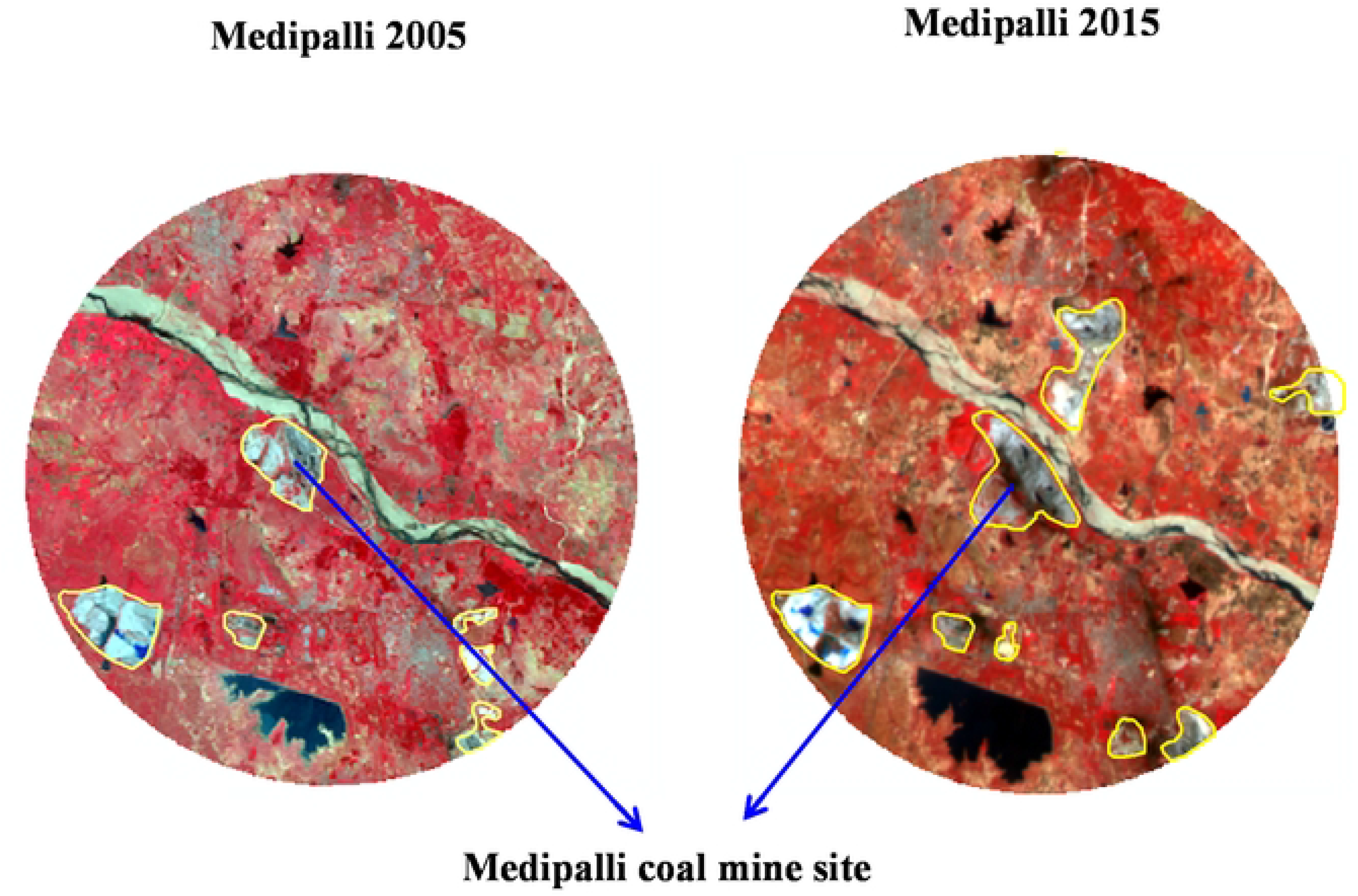
Satellite images of Medipalli open cast coal mine area a) Year 2005 b) Year 2015

**Figure 5.**
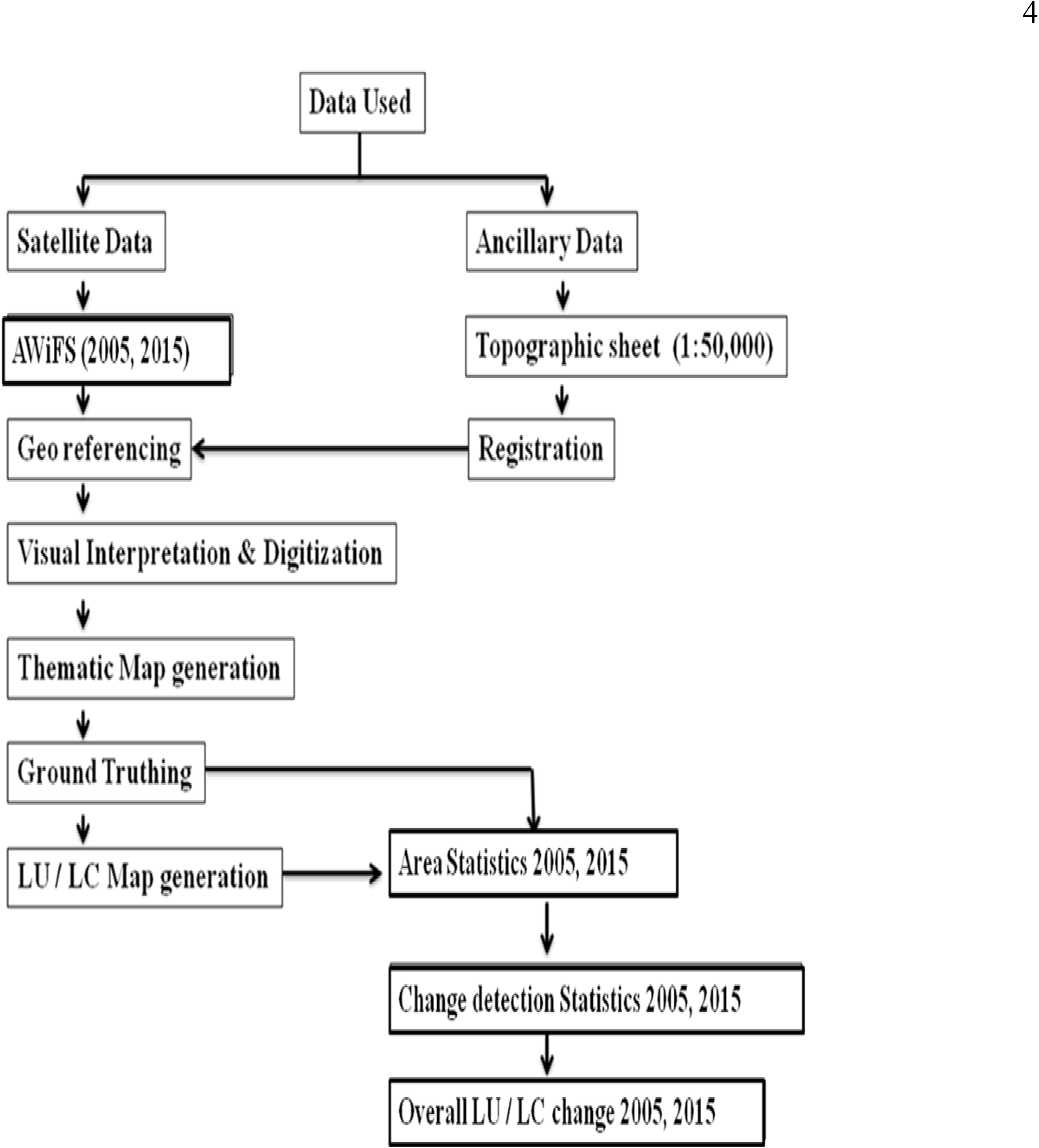
Flowchart demonstrating the methodology used to generate a land use/land cover map with corresponding change detection statistics.

The satellite photos of the Ramakrishnapur open cast coal mine region from 2005 to 2015 are presented in Fig 2, and the categorized map of the related images is shown in Figs 7 and 8, which illustrate eight categories of land use/land cover. Table 2 shows the area under various categories and dynamics in LU/LC, as well as the percentage area. Table 2 compares the extent of that LU/LC in 2005 and 2015. Positive values of % change indicate an increase, whereas negative values indicate a reduction. In Ramkrishnapur, mining was started just three years ago only, which was reflected in nil mining area during 2005. By the year 2015 it was increased to 1008 ha *i.e.,* 3.1 per cent which was depicted in pie diagram (Fig 6). This mine was only three years old when it began, therefore an average of 300 ha per year is being turned into mining territory. Mining activity increased during the course of the year, and enormous amounts of mining debris, such as overburden, were deposited on neighbouring land, damaging the quality of the soil and limiting its potential to produce, transforming it to barren land (41,42,43). Previously, this area was covered in forest, wetlands, and agricultural land. The removal of brush and agricultural areas during mining operations was often followed by significant environmental damage (18,19,44,50). According to the LU/LC change analysis, there was conversion and a decline in scrubland (sparse or open forest). When coal mine overburden is placed in regions that haven’t been mined, it produces mine spoils, which eventually have an impact on the nearby vegetation. Around the littoral swamp, the vegetation experienced a negative change of 9.7 %. The shallow water bodies had a decline of 9% during a ten-year period and were turned into mining areas (38,40), as seen plainly on satellite and classified maps. Table 2 makes it clear that built-up land has increased throughout this time, from 12.1% to 13.7% (Figs 4 and 5), including contributions from agricultural land and water bodies (32,33,45). 3.7% of water bodies were found to have changed negatively due to building up land.

**Figure 6.**
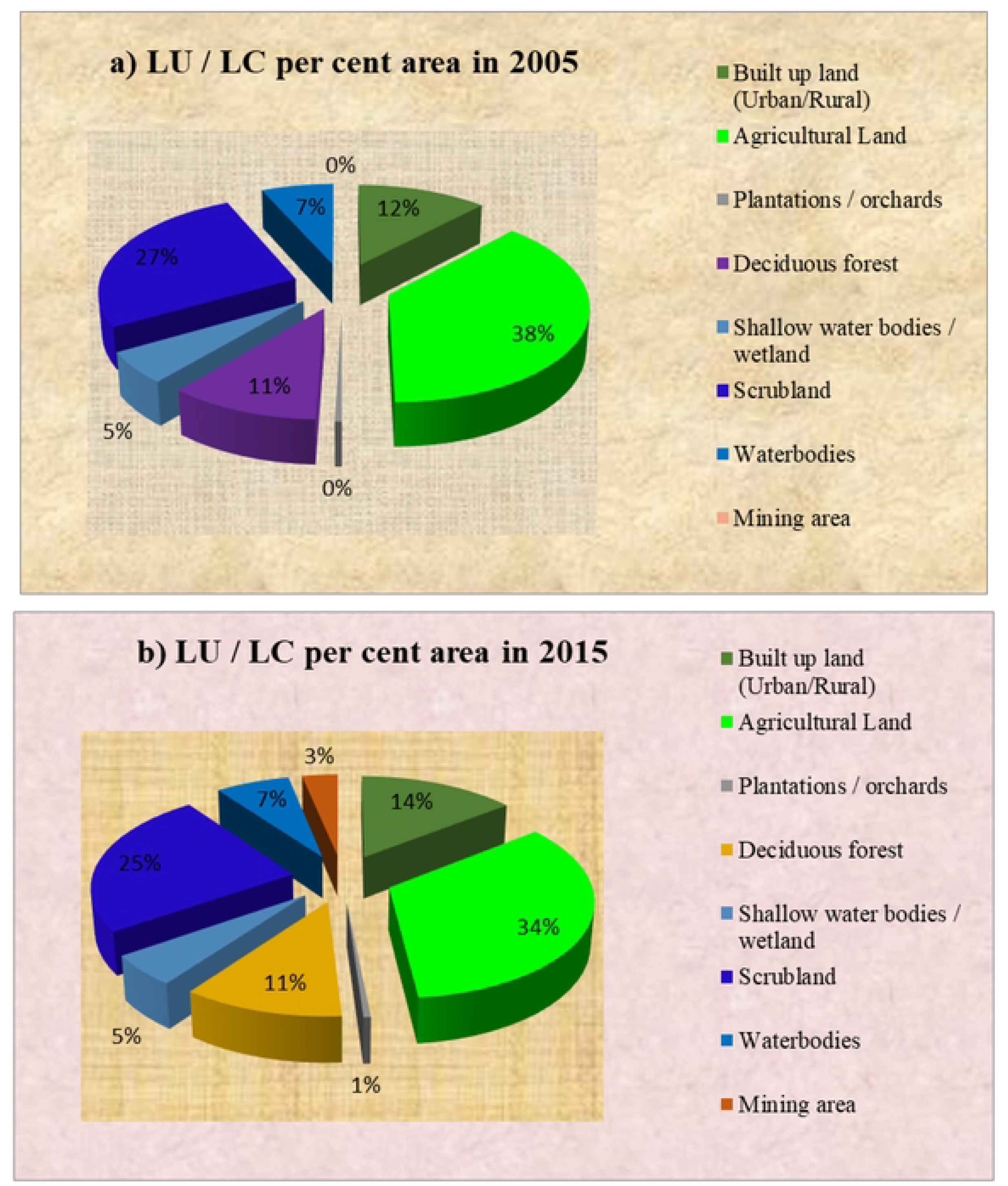
Pie chart showing the per cent area of Ramakrishnapur open cast coal mine a) year 2005 b) year 2015

**Figure 7.**
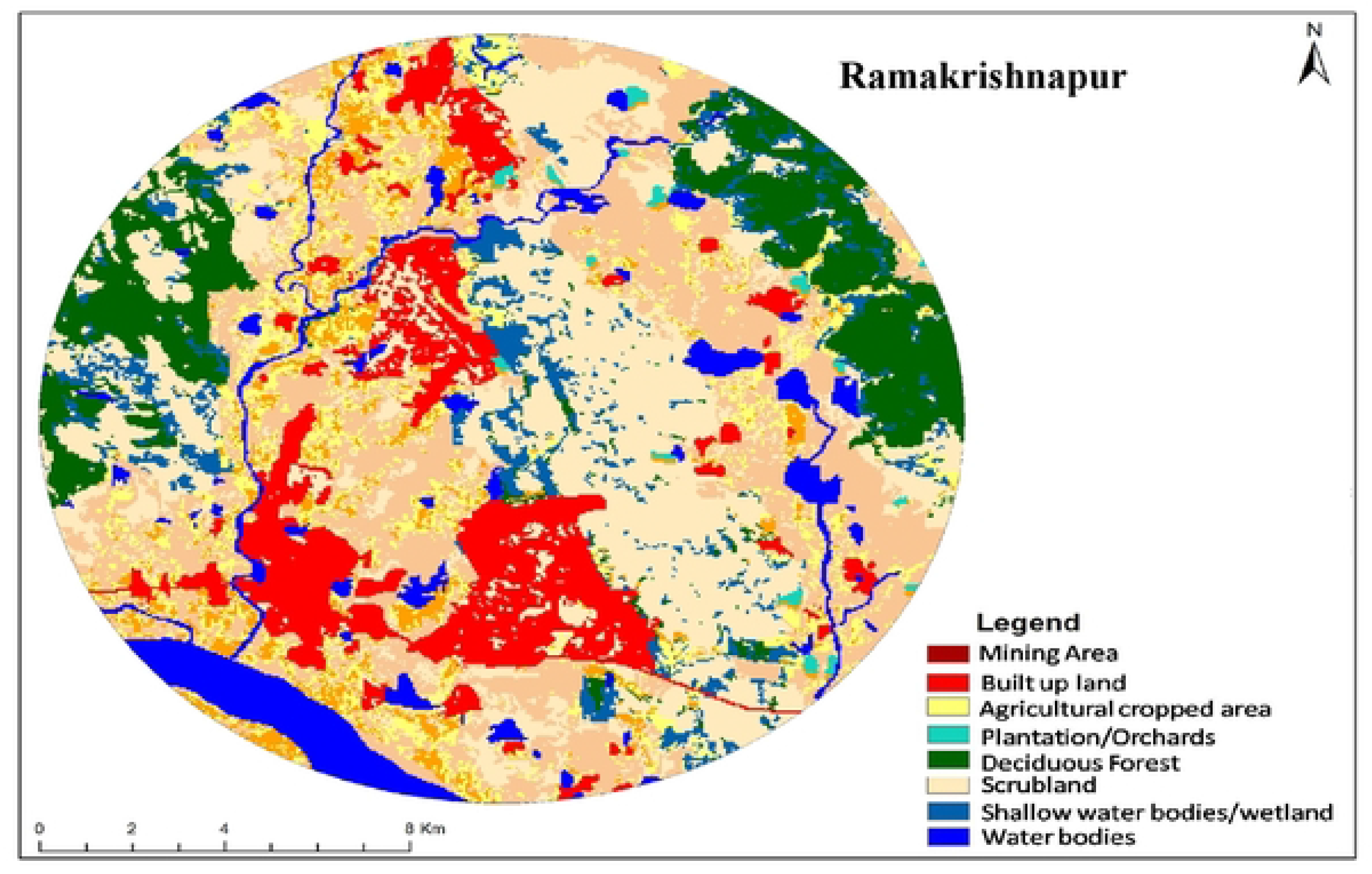
Land use *I* Land cover map of Ramakrishnapur open cast coal mine area (AWiFS, 2005)

**Figure 8.**
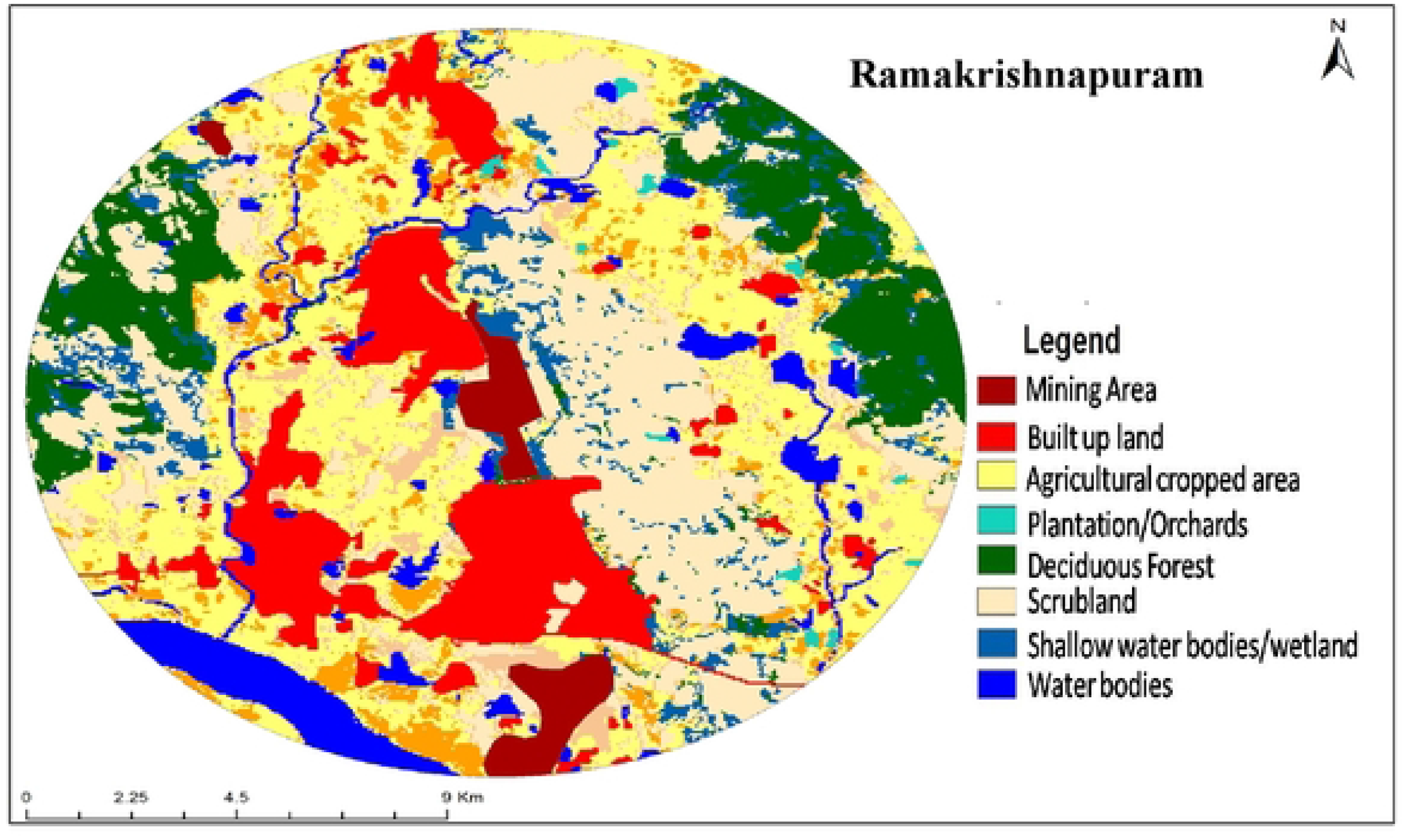
Raniakrishnapur open cast coal mine area land usage / land cover map (AWiFS, 2015)

**Table 2.**
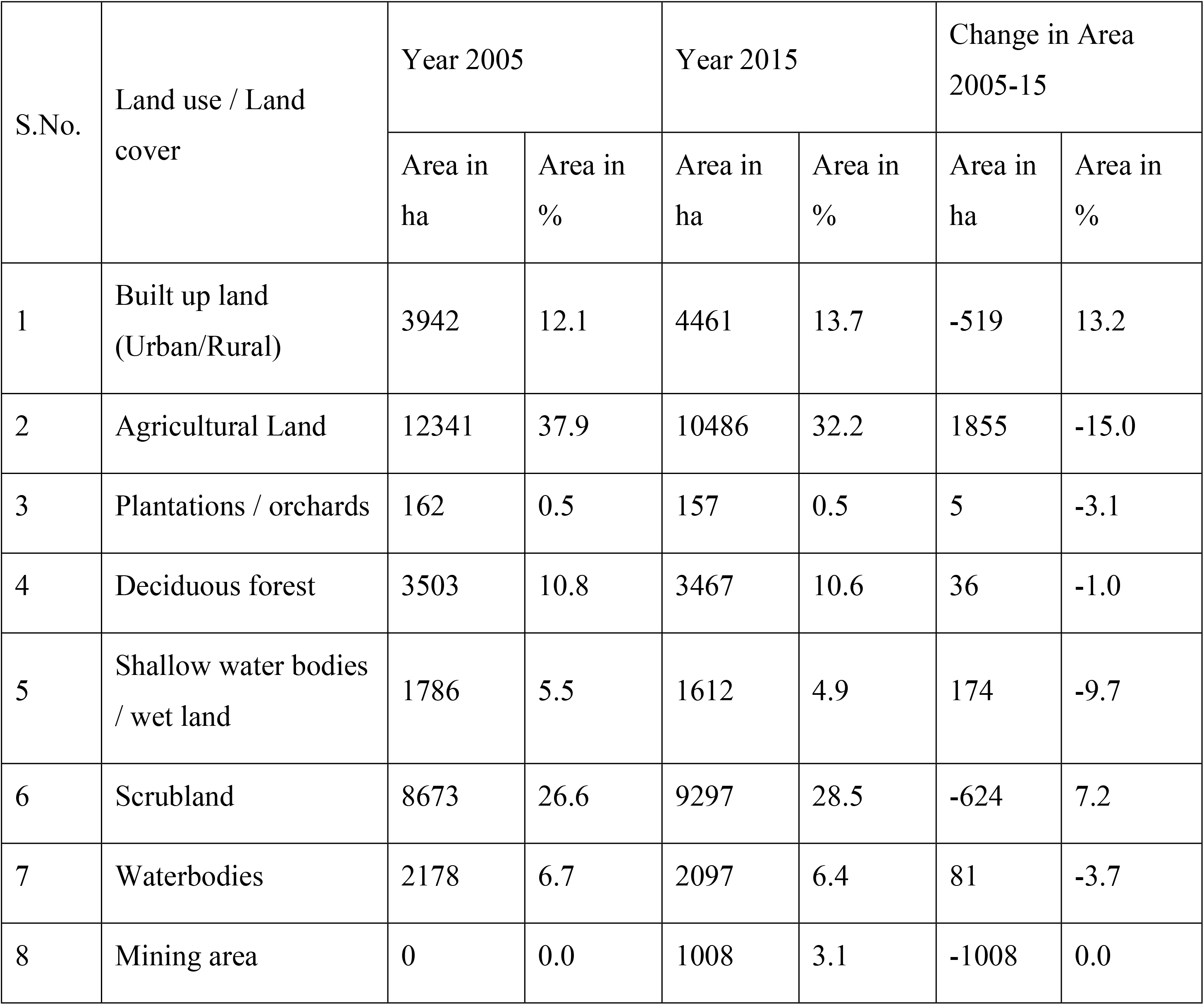
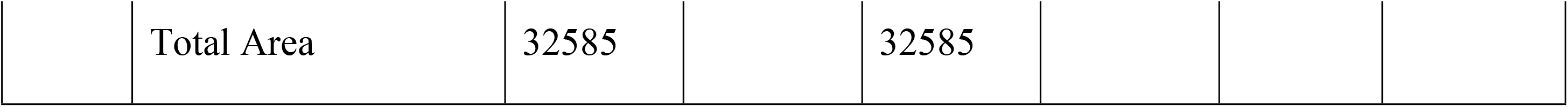
Land use / Land cover statistics and its dynamics during the study area from 2005-2015 at Ramakrishnapur open cast coal mine area.

The Satellite images of Srirampur open cast coal mine area of year 2005 and 2015 are shown in Fig 3 and classified map of the respective images are shown as Figs 10 and 11 which shows 8 categories of Land use/Land cover. The detailed percent change, area statistics and area dynamics are shown in Table 3. As this mine was started 10 years back *i.e.,* 2005, the mining area was 1.7 percent only (556 ha) which was increased to 6.3 per cent (2047 ha) which shows an average increase of 150 ha per year (Fig 9) the percent LU/LC change was 268.2 here the conversion of mining area was from Agriculture land, Scrub land and Shallow water body / wetlands which are associated with coal mines. The Deciduous Forest was decreased from 0.7 to 0.5 % in the area scrub land decreased from 23.2 to 18.8 %. Forest destruction during mining operations was always followed by severe environmental harm (35,36). The conversion of agricultural land, plantation land, and scrub land at this location also contributed to a favorable improvement in the built-up area (32,33). There was a small decline in water bodies of (0.3%), or from 9.9% to 9.6%, which indicates that there was reportedly no effect of coal mining on water bodies.

**Figure 9.**
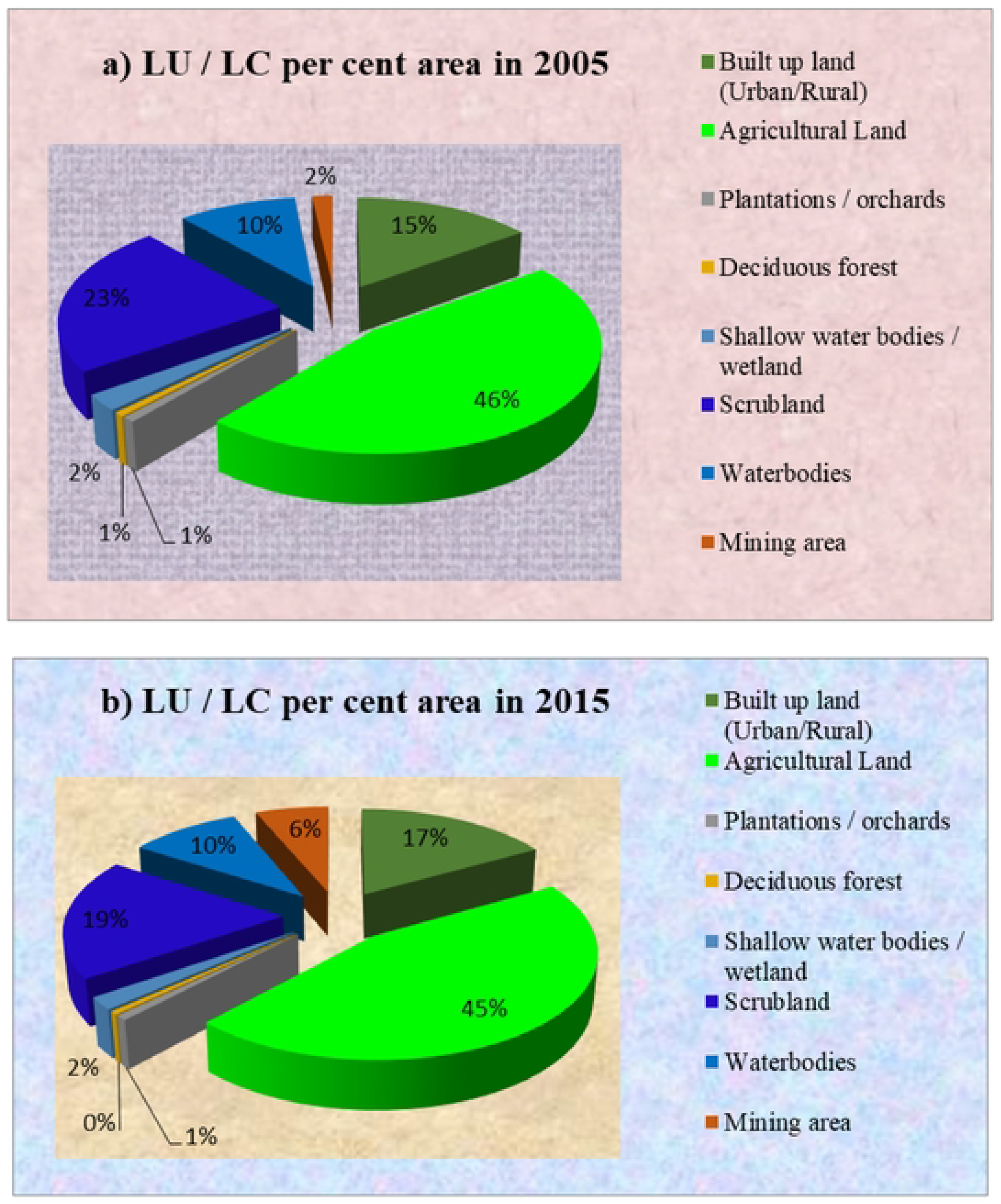
Pie chart showing the per cent area of Srirampur open cast coal mine a) year 2005 b) year 2015

**Figure 10.**
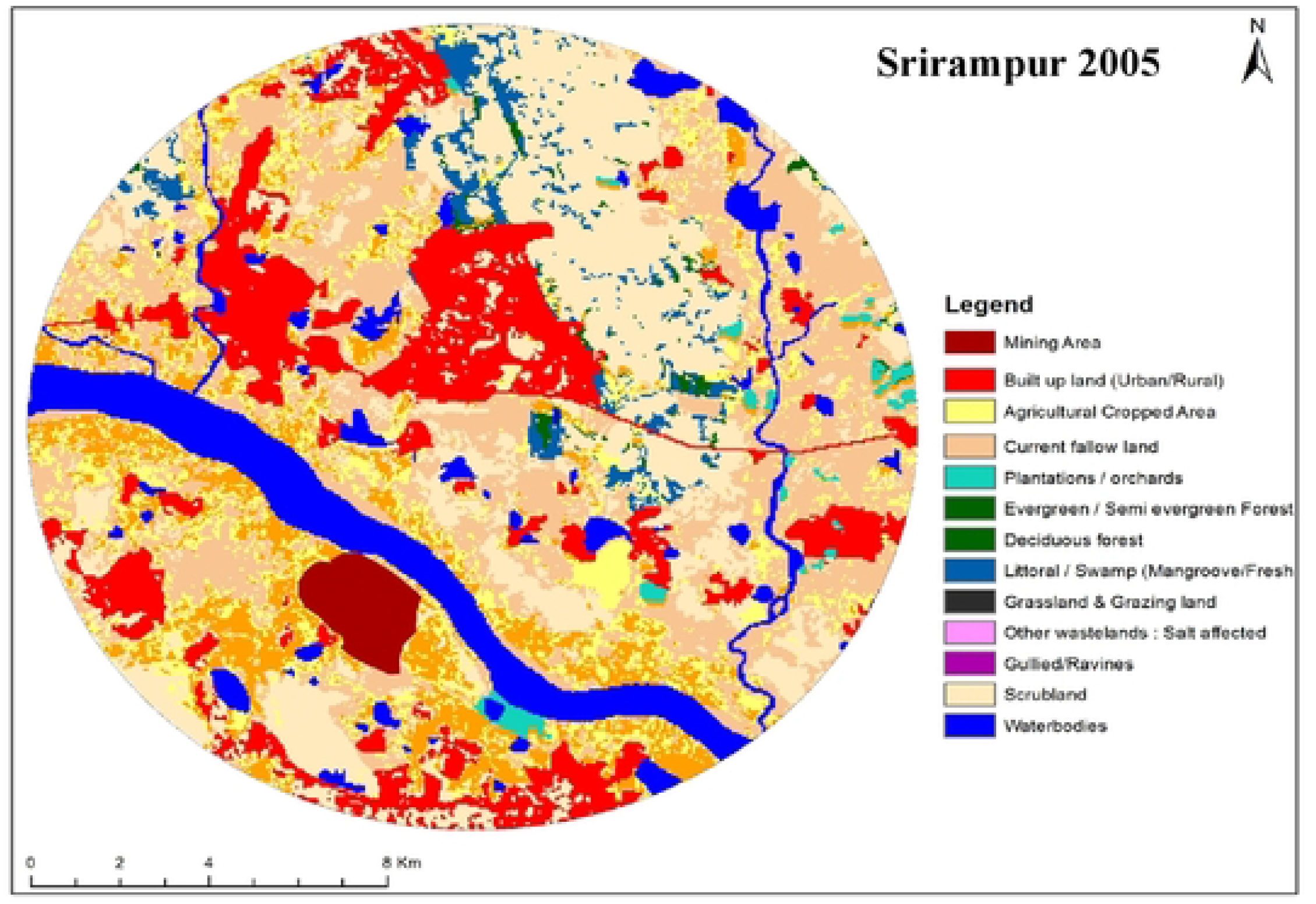
Land use / Land cover map of Srirampur open cast coal mine area (AWiFS, 2005)

**Figure 11.**
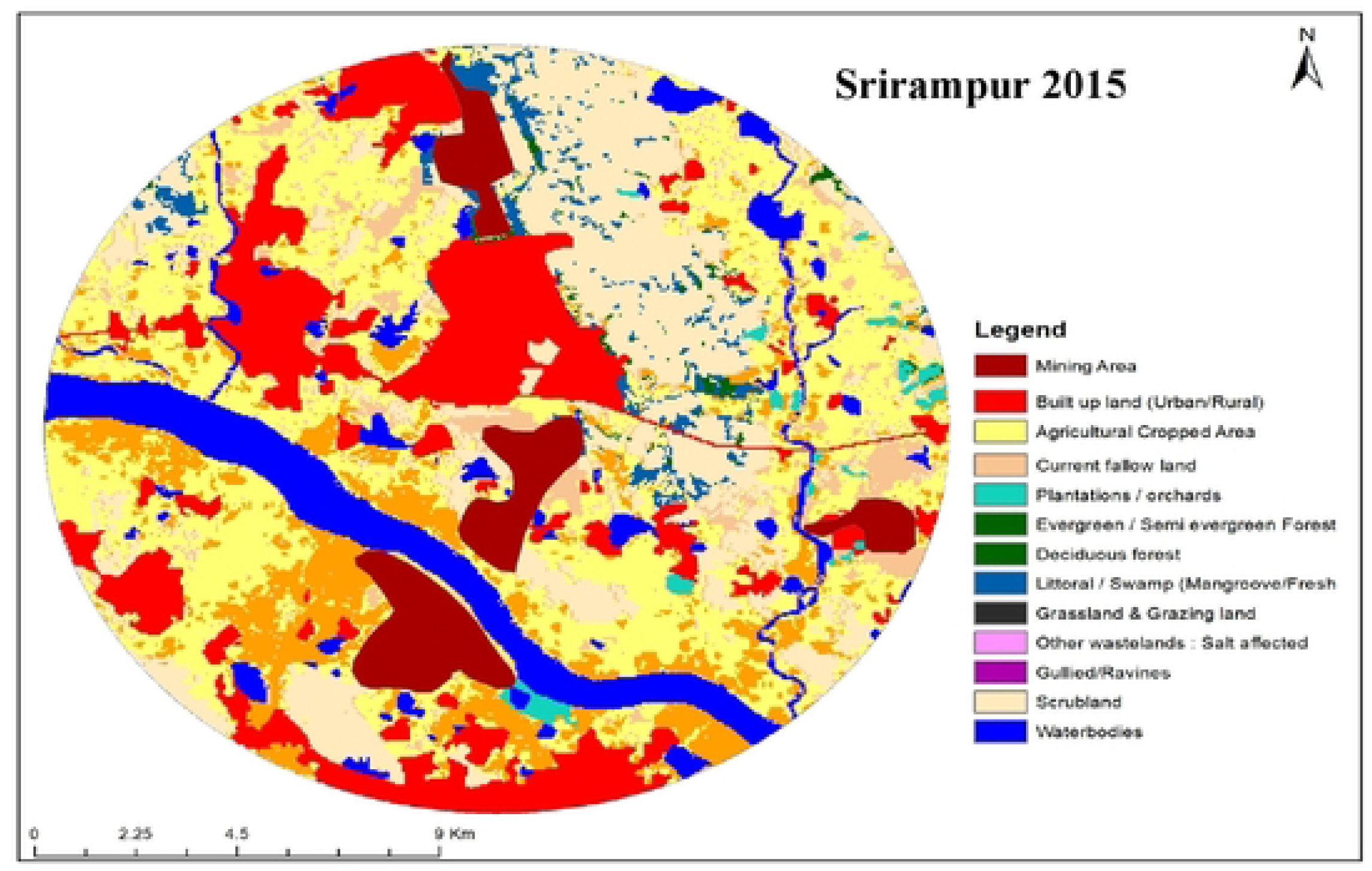
Land use / Land cover map of Srirampur open cast coal mine area (AWiFS, 2015)

**Table 3.**
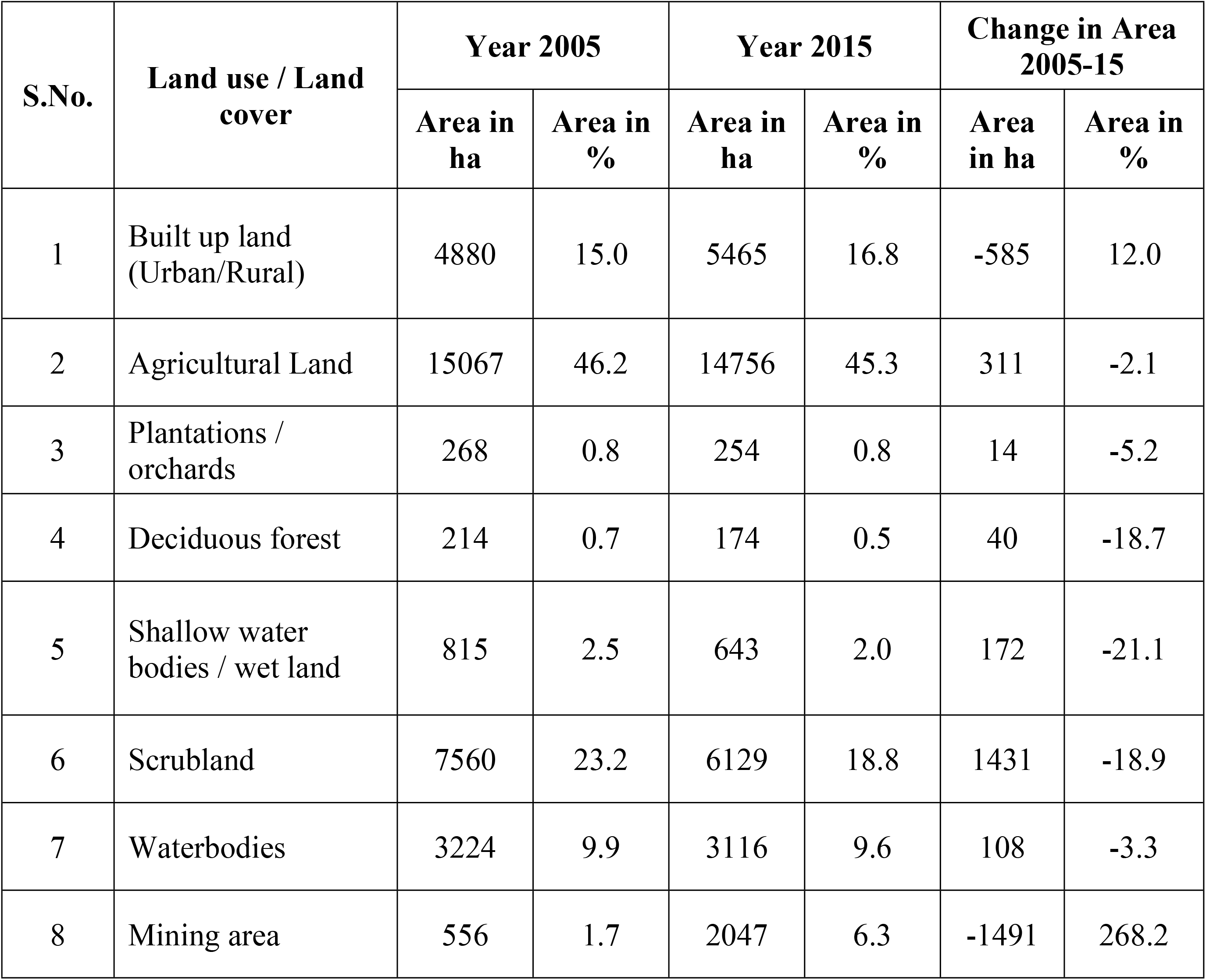
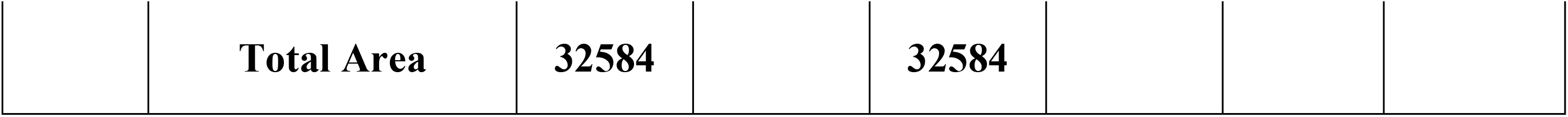
Land use / Land cover statistics and its dynamics during the study area from 2005-2015 at Srirampur open cast coal mine area.

The Satellite images of Medipalli open cast coal mine area of year 2005 and 2015 are shown in Fig 4 and classified map of the respective images are shown as Figs 13 and 14 which shows 8 categories of Land use/Land cover. The detailed percent change, area statistics and area dynamics are shown in Table 4. As this mining was started more than 15 years *i.e.,* 2005 the mining area was 4.9 percent only (1586 ha) which was increased to 7.9 per cent (2561 ha) which shows an average increase of 100 ha per year (Fig 12) the percent LU/LC change was 61.5 here the mining area is coming from Scrub land which is associated with coal mines. When mining operations destroyed scrub (sparse forest) forests, the ecosystem was almost always severely harmed (37).

**Figure 12.**
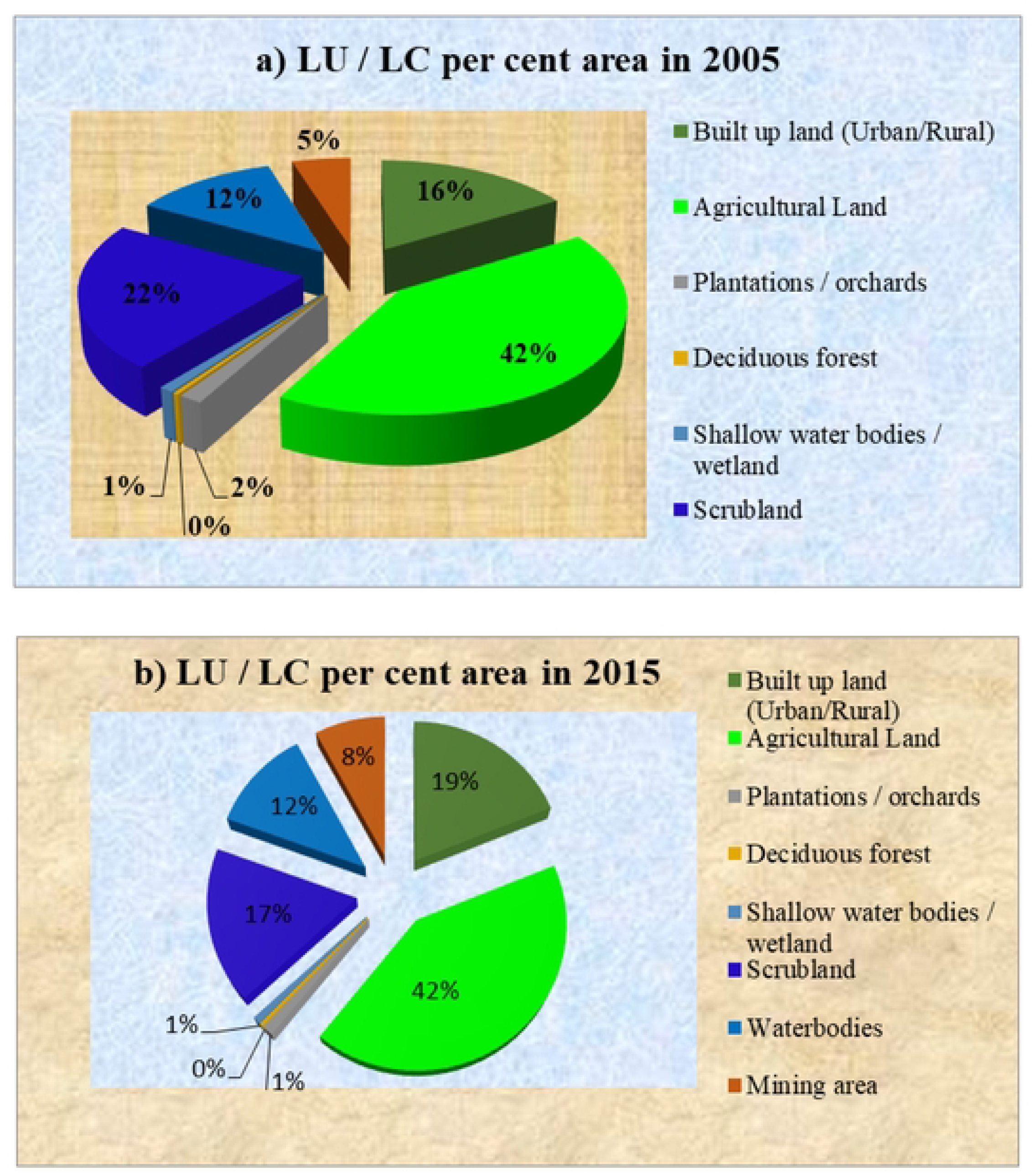
Pie chart showing the per cent area of IYIedipalli open cast coal mine a) year 2005 b) year 2015

**Figure 13.**
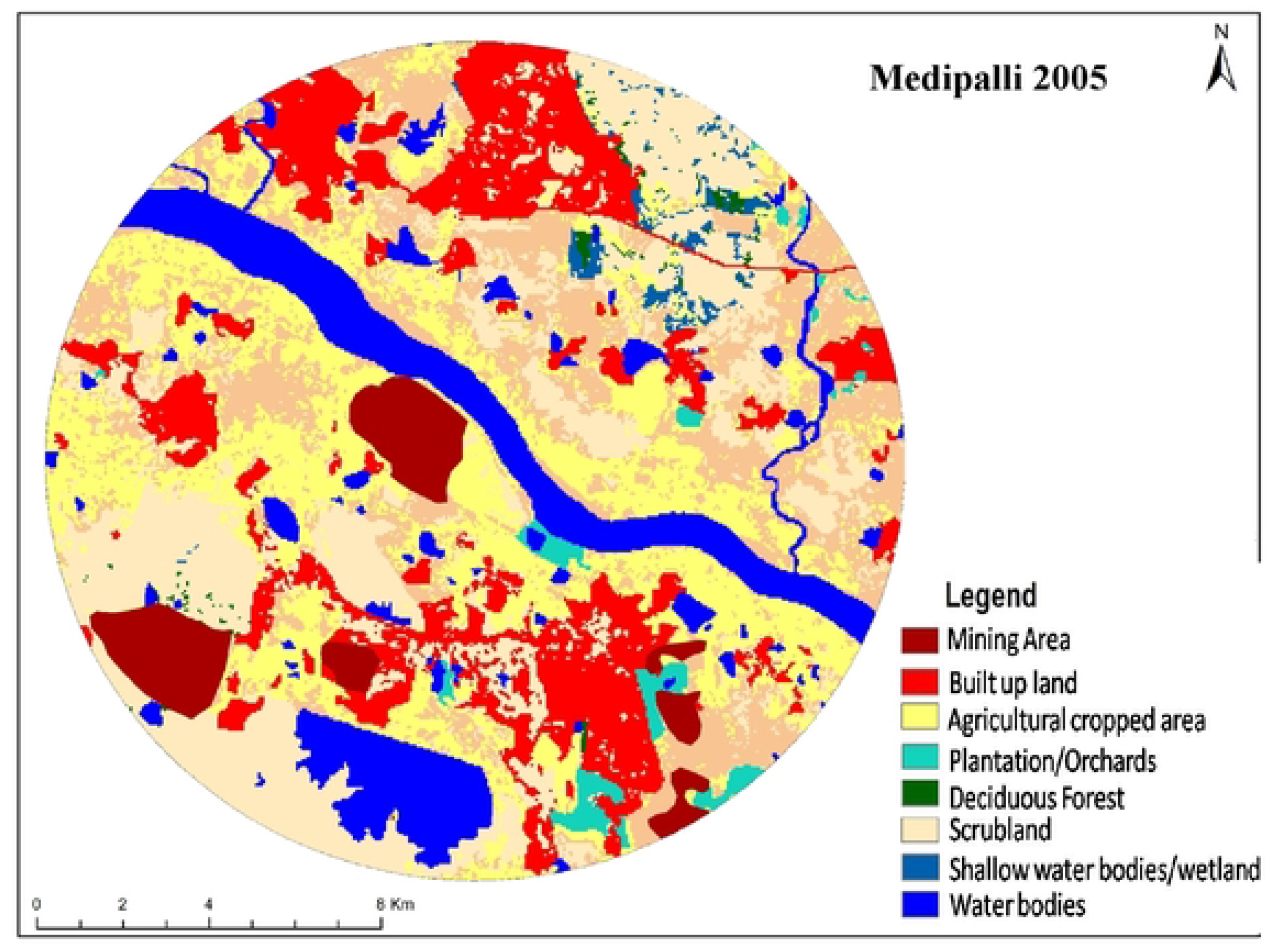
Land use / Land cover map of Medipalli open cast coal mine area (AWiFS, 2005)

**Figure 14.**
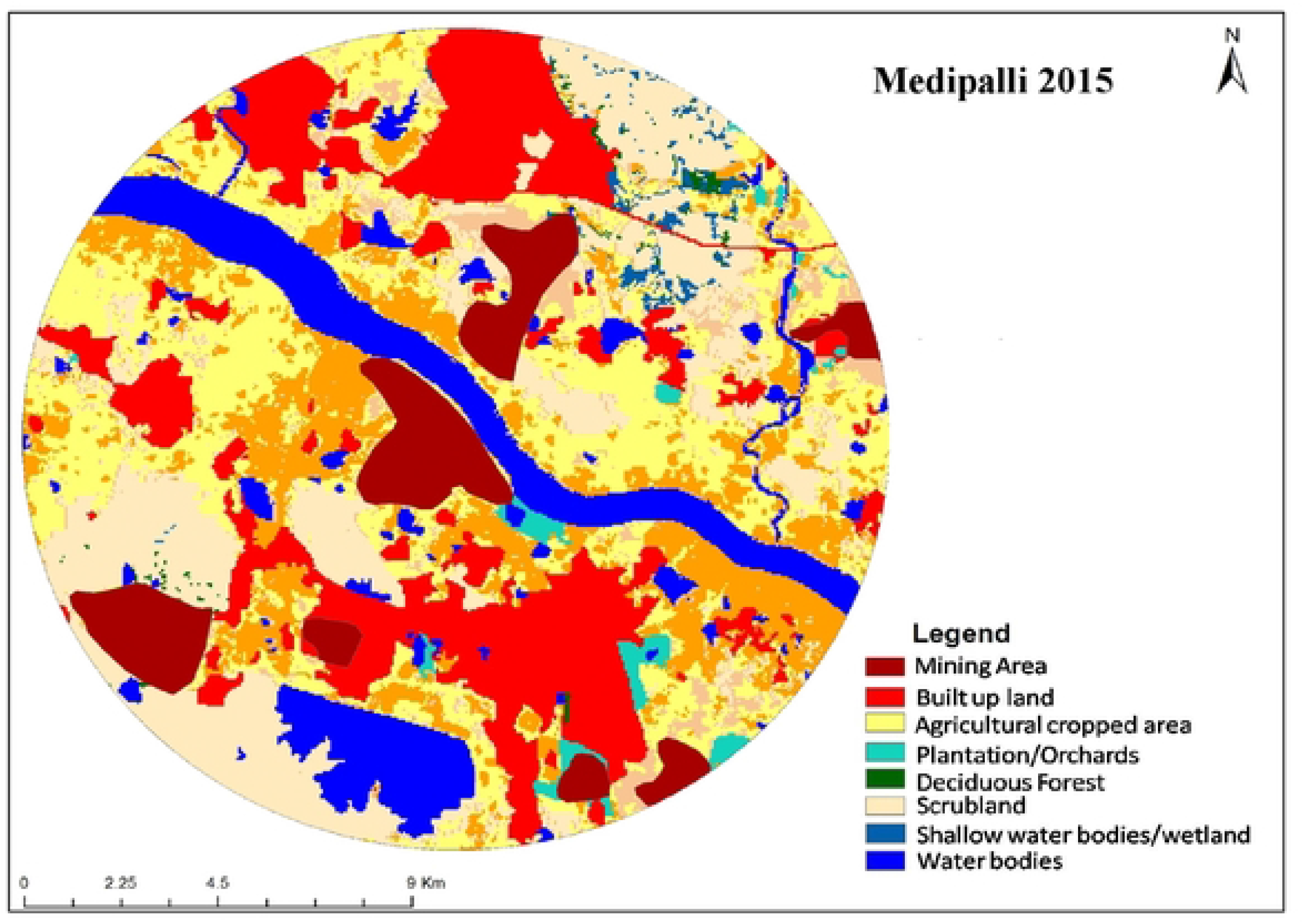
Land use / Land cover map of Medipalli open cast coal mine area (AWiFS, 2015)

**Table 4.**
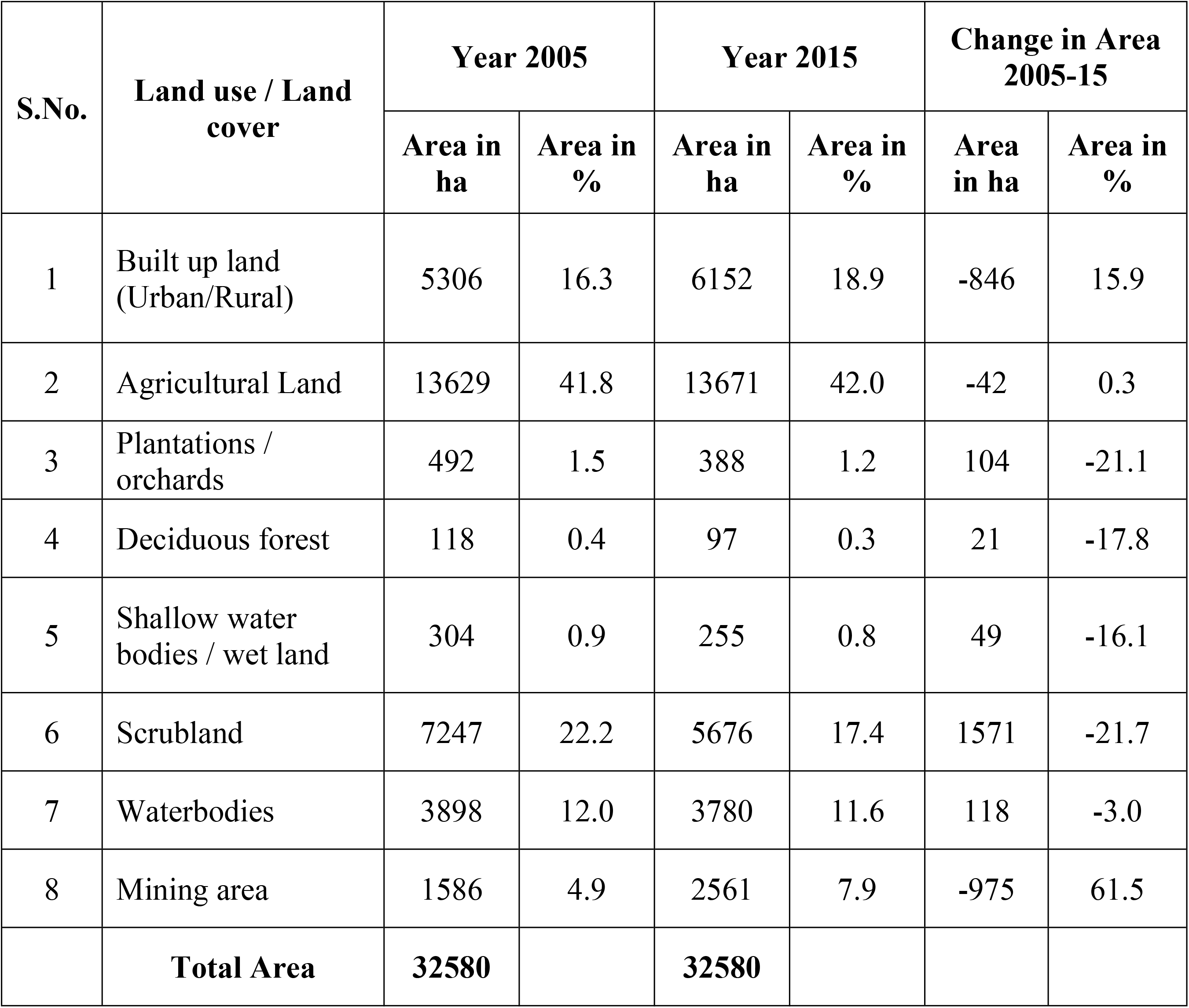
Land use / Land cover statistics and its dynamics during the study area from 2005-2015 at Medipalli open cast coal mine area.

On average, the extent of area converted into open cast coal mining per year was highest at Ramakrishnapur (@ 300 ha/year), followed by Srirampur (@ 150 ha/year), and Medipalli (@ 100 ha/year). Methods for remote sensing have proved their ability to generate reliable spatiotemporal data of LULC (14,1) and its variations in the coal mining area.

With the start of mining in Ramakrishnapur, there was a significant decline in agricultural land, shallow water bodies, and deciduous forests, indicating that these land uses had been converted to mining. In contrast, conversion of scrubland followed by shallow water bodies, agriculture areas, and deciduous forests into mining area was identified in the other two mining sites (Srirampur and Medipalli). However, the built-up land at all three mining sites came from cultivated areas, with scrubland also contributing at Medipalli and Srirampur. The overall impact of open cast mining on deep water bodies was not discovered. Long-term planning is critical for the sustainable and proper use of limited land and water resources in mining and adjacent regions (46,47,48,50).

## Conclusion

The extent of area converted into open cast coal mining per year was highest at Ramakrishnapur (@ 300 ha/year), followed by Srirampur (@ 150 ha/year), and Medipalli (@ 100 ha/year) as determined by the spatiotemporal statistics of LULC and its changes in coal mining area on average. With the start of mining, agricultural land, shallow water bodies, and deciduous forests in Ramakrishnapur significantly decreased; this was evidence of the change of these land uses to mining. The conversion of scrubland, followed by shallow water bodies, agricultural fields, and deciduous forests, into mining area was found at the other two mining sites (Srirampur and Medipalli). However, the developed land at each of the three mining sites came from cultivated land, with additional contributions from scrubland at Medipalli and Srirampur. On the whole, there was no evidence of an influence of open cast mining on deep water bodies. For the mining and neighbouring regions to manage the available land and water resources sustainably and effectively, careful long-term planning is essential.

## Acknowledgment

This research was funded and supported by the Professor Jayashankar Telangana State Agriculture University, Hyderabad, Telangana, India

## References

1. Jensen JR. Introductory digital image processing: A remote sensing perspective. 3rd ed. Upper Saddle River: Prentice Hall; 2005.

2. Lillesand TM, Kiefer RW, Chipman JW. Remote sensing and image interpretation. 7th ed. New York: Wiley; 2015.

3. Burrough PA, McDonnell RA. Principles of geographical information systems. Oxford: Oxford Univ Press; 1998.

4. Longley PA, Goodchild MF, Maguire DJ, Rhind DW. Geographic information systems and science. 4th ed. New York: Wiley; 2015.

5. Campbell JB, Wynne RH. Introduction to remote sensing. 5th ed. New York: Guilford Press; 2011.

6. Bandyopadhyay KK, Ghosh PK, Chaudhary RS, Maiti K, Mandal KG, Misra AK. Integrated nutrient management practices in soybean-based and sorghum in sole and intercropping system in a Vertisol. Indian J Agric Sci. 2004;74:55–63.

7. FAO. State of the world’s land and water resources. Rome: FAO; 2016.

8. Prakash A, Gupta RK. Land-use mapping and change detection in a coal mining area: a case study in the Jharia coal field, India. Int J Remote Sens. 1998;19(3):391–410.

9. Congalton RG, Green K. Assessing the accuracy of remotely sensed data. 2nd ed. Boca Raton: CRC Press; 2009.

10. Lu D, Mausel P, Brondizio E, Moran E. Change detection techniques. Int J Remote Sens. 2004;25(12):2365–2407.

11. Foody GM. Status of land cover classification accuracy assessment. Remote Sens Environ. 2002;80:185–201.

12. Song C, Woodcock CE, Seto KC, Lenney MP, Macomber SA. Classification accuracy issues in change analysis. Remote Sens Environ. 2001;75:230–244.

13. Chitade AZ, Katiyar SK. Colour-based image segmentation using k-means clustering. Int J Eng Sci Technol. 2010;2(10):5319–5325.

14. Singh A. Digital change detection techniques using remotely sensed data. Int J Remote Sens. 1989;10:989–1003.

15. Townsend PA, Helmers DP, Kingdon CC, McNeil BE, de Beurs KM, Eshleman KN. Changes in the extent of surface mining and reclamation. Environ Manage. 2009;44:1–15.

16. Demirel H, Anbarjafari G. Image resolution enhancement by using discrete and stationary wavelet decomposition. IEEE Trans Image Process. 2011;20(5):1458–1460.

17. Rathore CS, Wright R. Monitoring environmental impacts of surface coal mining. Int J Remote Sens. 1993;14:1021–1042.

18. Mishra VK, Rai PK, Mohan K. Evaluation of land use/land cover changes using remote sensing. Environ Monit Assess. 2010;165:193–203.

19. Pandey B, Agrawal M, Singh S. Assessment of air pollution around coal mining area. Environ Earth Sci. 2014;71:279–289.

20. Forman RTT. Land mosaics: The ecology of landscapes and regions. Cambridge: Cambridge Univ Press; 1995.

21. Turner MG, Gardner RH, O’Neill RV. Landscape ecology in theory and practice. New York: Springer; 2001.

22. Wu J. Landscape sustainability science. Landscape Ecol. 2013;28:999–1023.

23. ISRO. IRS Resourcesat data user handbook. Hyderabad: NRSC; 2011.

24. ERDAS. ERDAS Imagine user guide. Atlanta: Hexagon Geospatial; 2010.

25. Coppin P, Jonckheere I, Nackaerts K, Muys B, Lambin E. Digital change detection methods in ecosystem monitoring. Int J Remote Sens. 2004;25(9):1565–1596.

26. Belgiu M, Drăguţ L. Random forest in remote sensing: A review. ISPRS J Photogramm Remote Sens. 2016;114:24–31.

27. Maxwell AE, Warner TA, Fang F. Implementation of machine learning in remote sensing. Remote Sens. 2018;10:1–27.

28. Roy PS, Roy A, Joshi PK, Kale MP, Srivastava VK, Srivastava SK. Development of decadal land use database. Curr Sci. 2015;108(8):1430–1440.

29. NRSC. Land use land cover atlas of India. Hyderabad: NRSC; 2012.

30. Pontius RG Jr. Quantification error versus location error. Photogramm Eng Remote Sens. 2000;66:1011–1016.

31. Pontius RG Jr, Millones M. Death to Kappa. Landscape Ecol. 2011;26:133–146.

32. Foley JA, DeFries R, Asner GP, Barford C, Bonan G. Global consequences of land use. Science. 2005;309:570–574.

33. Lambin EF, Turner BL, Geist HJ, Agbola SB, Angelsen A. Land-use change drivers. Glob Environ Change. 2001;11:261–269.

34. Chavez PS. Image-based atmospheric corrections revisited. Photogramm Eng Remote Sens. 1996;62:1025–1036.

35. Sonter LJ, Moran CJ, Barrett DJ, Soares-Filho BS. Mining drives extensive deforestation in the Brazilian Amazon. Proc Natl Acad Sci USA. 2014;111:9899–9904.

36. Asner GP, Llactayo W, Tupayachi R, Luna ER. Elevated rates of gold mining in the Amazon. Proc Natl Acad Sci USA. 2009;106:206–211.

37. Ghose MK, Majee SR. Assessment of dust generation in mining areas. Environ Monit Assess. 2000;68:47–59.

38. McFeeters SK. The use of NDWI in the delineation of open water features. Int J Remote Sens. 1996;17:1425–1432.

39. Xu H. Modification of NDWI to enhance open water features. Int J Remote Sens. 2006;27:3025–3033.

40. Pekel JF, Cottam A, Gorelick N, Belward AS. High-resolution mapping of global surface water. Nature. 2016;540:418–422.

41. Ghose MK. Effect of opencast mining on soil fertility. Environ Monit Assess. 2004;97:43–54.

42. Brady NC, Weil RR. The nature and properties of soils. 15th ed. New York: Pearson; 2016.

43. Lal R. Soil carbon sequestration impacts on climate and food security. Science. 2004;304:1623–1627.

44. Areendran G, Raj K, Mazumdar S. Land use/land cover change analysis in mining areas. Ecol Indic. 2013;34:103–111.

45. Geist HJ, Lambin EF. Proximate causes of tropical deforestation. Bioscience. 2002;52:143–150.

46. IPCC. Climate change and land. Geneva: IPCC; 2019.

47. UNCCD. Global land outlook. Bonn: UNCCD; 2017.

48. Kumar A, Pandey AC. Evaluating impact of coal mining activity on land use/land cover using temporal satellite images in South Karanpura coalfields, Jharkhand, India. Int J Adv Remote Sens GIS. 2013;2(1):183–197.

49. Pal M, Mather PM. Support vector machines for classification in remote sensing. Int J Remote Sens. 2003;24:561–569.

50. Singh S, Srivastava PK, Singh D. Monitoring mining impact using geospatial techniques. J Indian Soc Remote Sens. 2017;45:875–885.

